# CIViC MCP: Integrating Large Language Models with the Clinical Interpretations of Variants in Cancer

**DOI:** 10.1101/2025.10.13.682185

**Authors:** Lars Schimmelpfennig, Quentin Cody, Joshua McMichael, Adam C. Coffman, Mariam Khanfar, Jinglun Li, Jennie Yao, Jason Saliba, Arpad Danos, Susanna Kiwala, Alex Wagner, Javier Sanz-Cruzado, Jake Lever, Malachi Griffith, Obi L. Griffith

## Abstract

**Summary:** The Clinical Interpretation of Variants in Cancer (CIViC) knowledgebase provides a community-driven, open-source platform for discussing the biological and clinical significance of molecular variants in cancer. To enable users to make complex connections between CIViC information, we developed the CIViC Model Context Protocol (MCP) server, which allows users to interface with the CIViC API through natural language via large language models (LLMs), facilitating the rapid summarization of expertly curated cancer variant interpretations.

**Availability and implementation:** The CIViC MCP server is detailed at https://github.com/griffithlab/civic-mcp-server. The repository includes instructions for accessing the server through the Claude desktop app (our recommended approach; Supplementary Figure 1) and hosting it locally with GPT-5, as well as a Python script for directly querying the MCP server. We also provide an MCP-supported Chatbot for CIViC users at https://civicdb.org/mcp-chat.

**Supplemental information:** Supplementary data are available at *Bioinformatics Advances* online.

## 1. Introduction

The identification of molecular variants drives precision oncology, informing clinical decision-making and treatment strategies. To give just one illustrative example, ALK fusion variants occur in fewer than 5% of non-small cell lung cancers but can be targeted by kinase inhibitors, such as crizotinib, for significant improvements in patient outcomes (Kwak *et al*. 2010). The complex mutational landscape of cancers makes the development of molecular variant knowledgebases critical to the goal of enabling the use of genomic information in precision oncology. To address this, the Clinical Interpretation of Variants in Cancer (CIViC) knowledgebase provides a community-driven, open-source platform for assessing the importance of molecular variants in cancer (available at civicdb.org) (Griffith *et al*. 2017; Danos *et al*. 2019; Krysiak et al. 2022). CIViC curates Evidence Items from peer-reviewed primary literature as its fundamental unit of information. Each item is assigned an Evidence Type that reflects the variant’s clinical or biological effect, including diagnostic, predictive, prognostic, predisposing, oncogenic, or functional. Each Evidence Type has a corresponding significance. For example, predictive evidence can indicate whether a tumor is sensitive or resistant to a specific therapy. CIViC Assertions are created to summarize collections of Evidence Items within a specific cancer context, adhering to recognized guidelines (Li *et al*. 2017; Horak et al. 2022).

**Figure 1.**
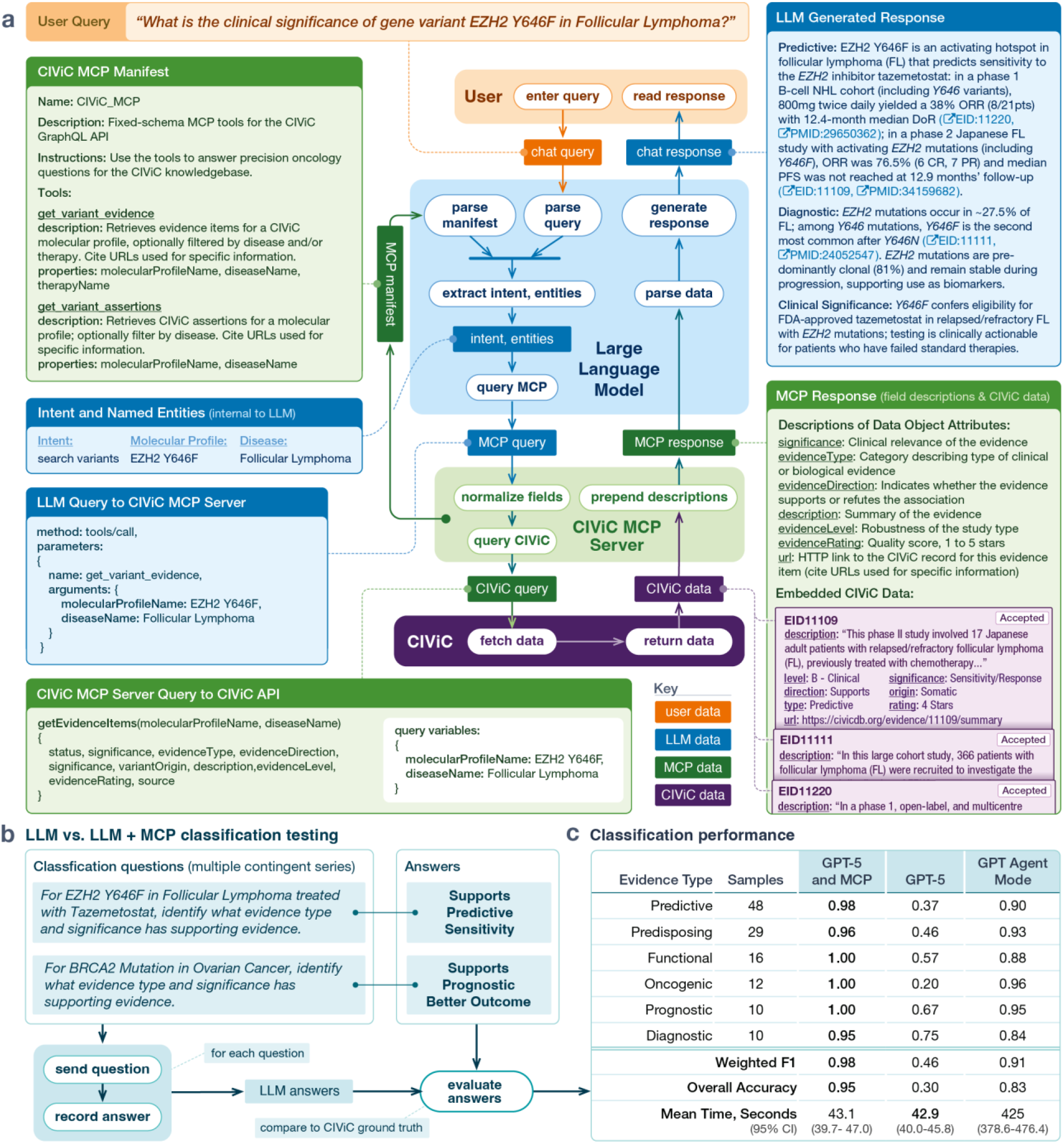
CIViC MCP workflow and evaluation. **(a) Workflow**. A user asks about the clinical significance of a variant-disease combination. The LLM reads the MCP manifest, extracts intent and named entities (molecular profile, disease, therapy), selects the appropriate tool, and issues a fixed-schema GraphQL query via the CIViC MCP server. The server queries the CIViC API, returns structured records enriched with field descriptions, and the LLM composes a cited summary. **(b) Evaluation setup**. For each of 100 CIViC triplets (molecular profile, disease, therapy), the model was given a fixed list of evidence type-significance pairs and asked to classify each as supported, not supported, both, or lacking evidence based on the direction of matching CIViC records. **(c) Classification performance**. GPT-5 + MCP achieves the highest weighted F1 across all Evidence Types while maintaining comparable response times to GPT-5 alone. Agent Mode narrows the accuracy gap but at roughly 10-fold greater latency.

LLMs provide a natural-language interface to CIViC, enabling users to discover, aggregate, and summarize curated oncology knowledge without navigating to multiple pages. They excel at transforming unstructured biomedical text into structured outputs through few-or zero-shot learning and entity normalization. However, LLMs cannot guarantee coverage of specialized, rapidly updated resources like CIViC from pretraining alone, and they may misinterpret CIViC’s data model, fabricating details or citations when asked for fine-grained clinical information (Bhattacharyya et al. 2023). Without API integration, chatbots typically fall back on search-based web retrieval, identifying relevant pages through a search engine and then reading their contents. As a result, discoverability is constrained by the effectiveness of the search mechanism. Agentic web or UI interaction offers a different form of indirect access, in which the model navigates websites by simulating human inputs. However, for a highly structured resource such as CIViC, it remains unclear whether this approach can match the accuracy, provenance, and efficiency of direct structured retrieval. In this work, we evaluate whether direct, structured API access through a standardized protocol can close this gap, improving both accuracy and efficiency over unassisted and agent-based approaches. Additionally, because CIViC’s curation is intentionally granular, captured in individual evidence items for transparency and provenance, many users may benefit from a conversational layer that retrieves the right items and composes succinct, well-cited answers. To provide this layer, we developed a Model Context Protocol (MCP) server that allows LLMs to query CIViC information seamlessly and provide detailed answers to users (Figure 1). MCP servers expose external tools and data sources through a simple, standardized interface, enabling LLMs to issue user-guided queries and receive structured results. In biomedicine, MCP servers are increasingly wrapping domain APIs, databases, and literature search endpoints (Kuehl *et al*. 2025). The CIViC MCP server benefits both new and experienced users by supporting complex, reproducible queries and rapid, citation-rich summaries.

The CIViC MCP server leverages CIViC’s public GraphQL API to facilitate user-guided queries. The GraphQL schema makes the API straightforward for LLMs to use as objects are explicitly linked (e.g., Evidence Item → Molecular Profile → Variant), letting models see how to move from a clinical question to the exact fields it needs. Through introspection, the LLM reads the schema, discovers types, fields, arguments, and deprecations, and then composes precise queries that fetch only the necessary data. The CIViC knowledge model is highly domain-specific, including complex concepts such as “Molecular Profiles” and “Assertions,” which have precise meanings specific to the platform and may be misinterpreted by an LLM (Krysiak *et al*. 2022). Despite GraphQL introspection, early tests revealed that Claude Sonnet 4 frequently mis-specified types and arguments, leading to invalid or overly broad queries. While introspection did improve reliability on later queries, where the LLM could contextualize GraphQL error messages against the schema and refine its requests, it did not close the gap relative to predefined tools. We circumvent this challenge in the current implementation of the CIViC MCP server by predefining the GraphQL queries, rather than allowing an LLM to write them to ensure reliability. We note that these challenges are not specific to GraphQL. A REST API would similarly require the LLM to know the correct endpoints, parameter names, and expected values, information that is available in documentation but that current LLMs cannot reliably operationalize without a structured tool interface.

## 2. CIViC MCP Server

The CIViC MCP server is hosted on Cloudflare and is publicly accessible. It provides two tools based on predefined GraphQL queries: one for retrieving CIViC Evidence Items and one for retrieving CIViC Assertions. For both tools, a Molecular Profile name is required, while cancer type and therapy may be provided as optional filters to further narrow the results. The Evidence Item tool retrieves curated information specific to individual studies, including an assessment of evidence strength, a summary of study findings within the relevant clinical context, and associated PubMed IDs. The Assertion tool retrieves higher-level summaries that synthesize collections of Evidence Items within a specific cancer-variant context. These tools were made separate to accommodate questions requiring different levels of granularity. Each response includes the relevant CIViC URLs, enabling direct verification of CIViC-specific claims generated by the LLM. Evidence Item responses also include the associated PubMed ID for validation against the source literature. To support accurate summarization, the MCP server additionally returns definitions for each CIViC field.

To handle spelling differences and alternate names, the MCP server normalizes user inputs (molecular profiles, cancer types, therapies) to the closest CIViC preferred label. The preferred label is the canonical, API-recognized name used for exact matching. This normalization relies on precomputed alias lists that were generated from external and CIViC-curated resources, rather than querying those resources at runtime. For gene-level aliases, we used the VICC Gene Normalization Service to assemble synonym lists for genes appearing in CIViC molecular profiles (Kuzma, Stevenson and Wagner 2025). Variant aliases were obtained from CIViC’s curated alias fields. Cancer type aliases were derived from the Disease Ontology, and therapy aliases were derived from the NCI Thesaurus (Sioutos *et al*. 2007; Schriml *et al*. 2012). The MCP server scores candidates with Dice’s coefficient over character bigrams and selects the highest-scoring alias. We chose the Dice-Sørensen coefficient because it is fast and memory-light, making it a good fit for Cloudflare Workers despite more complex matchers offering higher accuracy.

### 2.1 Example Use Case

A user asks, “What is the clinical significance of EZH2 Y646F in Follicular Lymphoma?” (Figure 1a). The LLM extracts the entities (molecular profile, disease, therapy if present) and normalizes them to CIViC preferred labels (in this case, molecular profile is “EZH2 Y646F” and disease is “Follicular Lymphoma”). It selects the get_variant_evidence tool and supplies the necessary parameters. The MCP server then issues a predefined GraphQL query to the CIViC API and returns structured records: Evidence Items with fields such as evidence type, significance, direction, disease, therapy, description, evidence level, rating, and URLs. The LLM then uses only this returned CIViC content to compose an information-rich answer, linking back to the specific CIViC entries and source publications provided in the MCP response. In this case, it summarizes the clinical relevance, including describing several clinical trials, for EZH2 Y646F’s role as an activating mutation in follicular lymphoma (FL), a potential diagnostic biomarker for FL, and a biomarker of sensitivity to tazemetostat.

## 3. Benchmarking Accuracy and Efficiency of LLM Retrieval Methods for CIViC

We selected a controlled classification task: given a CIViC triplet (molecular profile, cancer type, therapy), the LLM must determine which CIViC clinical significance has supporting evidence. To assess how effectively LLMs can retrieve and interpret current CIViC information through the MCP server, we compared GPT-5’s performance with and without the MCP server on the same set of 100 randomly selected triplets from CIViC Evidence Items. We used a zero-shot prompt at temperature = 1 (Supplementary Prompt 1). We prompted GPT-5 to determine, for each candidate CIViC clinical significance, whether the available evidence supported that significance, did not support it, or provided no evidence for it. These three outcomes served as the class labels for evaluation, allowing us to distinguish between positive evidence, contradictory evidence, and absence of evidence. In CIViC, significance labels are grouped within Evidence Types, which broadly indicate the type of clinical or biological association described by an Evidence Item’s clinical summary. Each triplet could have multiple associated Evidence Types and significances. Performance was evaluated at the significance-label level and then summarized at the level of each Evidence Type by micro-averaging across all associated significance labels within that type (Figure 1b). We excluded the common “No Evidence” label to avoid inflating global metrics and obscuring errors. GPT-5 with MCP substantially outperformed GPT-5 alone (Figure 1c). GPT-5 + MCP achieved 0.95 overall accuracy and a 0.98 weighted F1, compared with 0.30 overall accuracy and 0.46 weighted F1 for GPT-5 without MCP. We also evaluated GPT Agent Mode, which interacts with websites through simulated browsing. Additional details of the Agent Mode benchmarking procedure are provided in the Supplementary Methods. Agent Mode improved markedly over GPT-5 alone, achieving 0.83 overall accuracy and a 0.91 weighted F1, but remained below GPT-5 + MCP while requiring nearly 10-fold longer runtime. Mean response times were similar for GPT-5 + MCP (43.1 s, 95% CI: 39.7-47.0) and GPT-5 alone (42.9 s, 95% CI: 40.0-45.8), whereas Agent Mode required 425.0 s (95% CI: 378.6-476.4). Together, these results show that MCP-based access to CIViC substantially improves classification performance without adding measurable latency.

## 4. Conclusion

We demonstrate that MCP servers provide standardized, LLM-friendly gateways to curated biomedical databases, enabling precise retrieval and structured summarization. Connecting CIViC to LLMs through MCP improves the accuracy of extracting clinical significance in variant-disease-therapy contexts compared with using an LLM’s web client, and it reduces latency relative to agent-mode browsing while maintaining fidelity. By offering structured, real-time access to curated oncology knowledge, the CIViC MCP server mitigates gaps in pretraining coverage, reduces hallucination risk, and yields reproducible, citation-rich outputs. MCP-mediated access both improves performance and shortens time to answer. Future work will develop constrained query-composition methods that complement predefined tools, enabling the safe execution of LLM-written queries to address unanticipated user questions. The CIViC MCP server can also be extended to integrate with complementary knowledgebases (e.g., DGIdb, ClinVar, OncoKB), enabling unified multi-database retrieval and cross-knowledge summarization through a single conversational interface.

## Author Contributions

Lars Schimmelpfennig (Formal analysis [lead], Investigation [lead], Writing-original draft [lead], Software [lead]), Quentin Cody, (Software [lead: created the original MCP server], Writing-review & editing [supporting]), Joshua McMichael (Visualization [lead]), Adam C. Coffman (Software [lead: created the CIViC MCP Chatbot], Writing-review & editing [supporting]), Mariam Khanfar (Investigation [supporting]), Jinglun Li (Investigation [supporting]), Jennie Yao (Investigation [supporting]), Jason Saliba (Writing-review & editing [supporting]), Arpad Danos (Writing-review & editing [supporting]), Susanna Kiwala (Writing-review & editing [supporting]), Alex H. Wagner (Writing-review & editing [supporting]), Javier Sanz-Cruzado (Writing-review & editing [supporting]), Jake Lever (Writing-review & editing [supporting], Supervision [lead]), Malachi Griffith (Writing-review & editing [supporting], Supervision [lead]), Obi L. Griffith (Writing-review & editing [supporting], Supervision [lead]).

## Conflict of interest

None declared.

## Funding

This work was supported by the National Cancer Institute (NCI) of NIH award numbers U24CA305456, U24CA237719, U24CA258115, and U24CA275783, and by the National Center for Advancing Translational Sciences (NCATS) of NIH under award number UL1TR002345 (Institute of Clinical and Translational Sciences [ICTS]).

## Notes

https://github.com/griffithlab/civic-mcp-server

### Competing Interest Statement

The authors have declared no competing interest.

### Summary of Updates

Evaluation section and figure was updated. Prompting strategy was redesigned to allow for running GPT Agent Mode across all samples for a stronger comparison.

## References

Bhattacharyya M, Miller VM, Bhattacharyya D et al. High Rates of Fabricated and Inaccurate References in ChatGPT-Generated Medical Content. Cureus 15:e39238.

Danos AM, Krysiak K, Barnell EK et al. Standard operating procedure for curation and clinical interpretation of variants in cancer. Genome Med 2019;11:76.

Griffith M, Spies NC, Krysiak K et al. CIViC is a community knowledgebase for expert crowdsourcing the clinical interpretation of variants in cancer. Nat Genet 2017;49:170–4.

Horak P, Griffith M, Danos AM et al. Standards for the classification of pathogenicity of somatic variants in cancer (oncogenicity): Joint recommendations of Clinical Genome Resource (ClinGen), Cancer Genomics Consortium (CGC), and Variant Interpretation for Cancer Consortium (VICC). Genet Med Off J Am Coll Med Genet 2022;24:986–98.

Koh JY, Lo R, Jang L et al. VisualWebArena: Evaluating Multimodal Agents on Realistic Visual Web Tasks. In: Ku L-W, Martins A, Srikumar V (eds.), Proceedings of the 62nd Annual Meeting of the Association for Computational Linguistics (Volume 1: Long Papers). Bangkok, Thailand: Association for Computational Linguistics, 2024, 881–905.

Krysiak K, Danos AM, Saliba J et al. CIViCdb 2022: evolution of an open-access cancer variant interpretation knowledgebase. Nucleic Acids Res 2022;51:D1230–41.

Kuehl M, Schaub DP, Carli F et al. Community-based biomedical context to unlock agentic systems. 2025:2025.07.21.665729.

Kuzma K, Stevenson J, Wagner A. VICC Gene Normalization Service. 2025, DOI: 10.5281/zenodo.16272753.

Kwak EL, Bang Y-J, Camidge DR et al. Anaplastic lymphoma kinase inhibition in non-small-cell lung cancer. N Engl J Med 2010;363:1693–703.

Li MM, Datto M, Duncavage EJ et al. Standards and Guidelines for the Interpretation and Reporting of Sequence Variants in Cancer. J Mol Diagn JMD 2017;19:4–23.

Schriml LM, Arze C, Nadendla S et al. Disease Ontology: a backbone for disease semantic integration. Nucleic Acids Res 2012;40:D940–6.

Sioutos N, de Coronado S, Haber MW et al. NCI Thesaurus: a semantic model integrating cancer-related clinical and molecular information. J Biomed Inform 2007;40:30–43.

Yoran O, Amouyal SJ, Malaviya C et al. AssistantBench: Can Web Agents Solve Realistic and Time-Consuming Tasks? In: Al-Onaizan Y, Bansal M, Chen Y-N (eds.), Proceedings of the 2024 Conference on Empirical Methods in Natural Language Processing. Miami, Florida, USA: Association for Computational Linguistics, 2024, 8938–68.

