## Supplemental Materials for "CIViC MCP: Integrating Large Language Models with the Clinical Interpretations of Variants in Cancer"

**Supplementary Data**

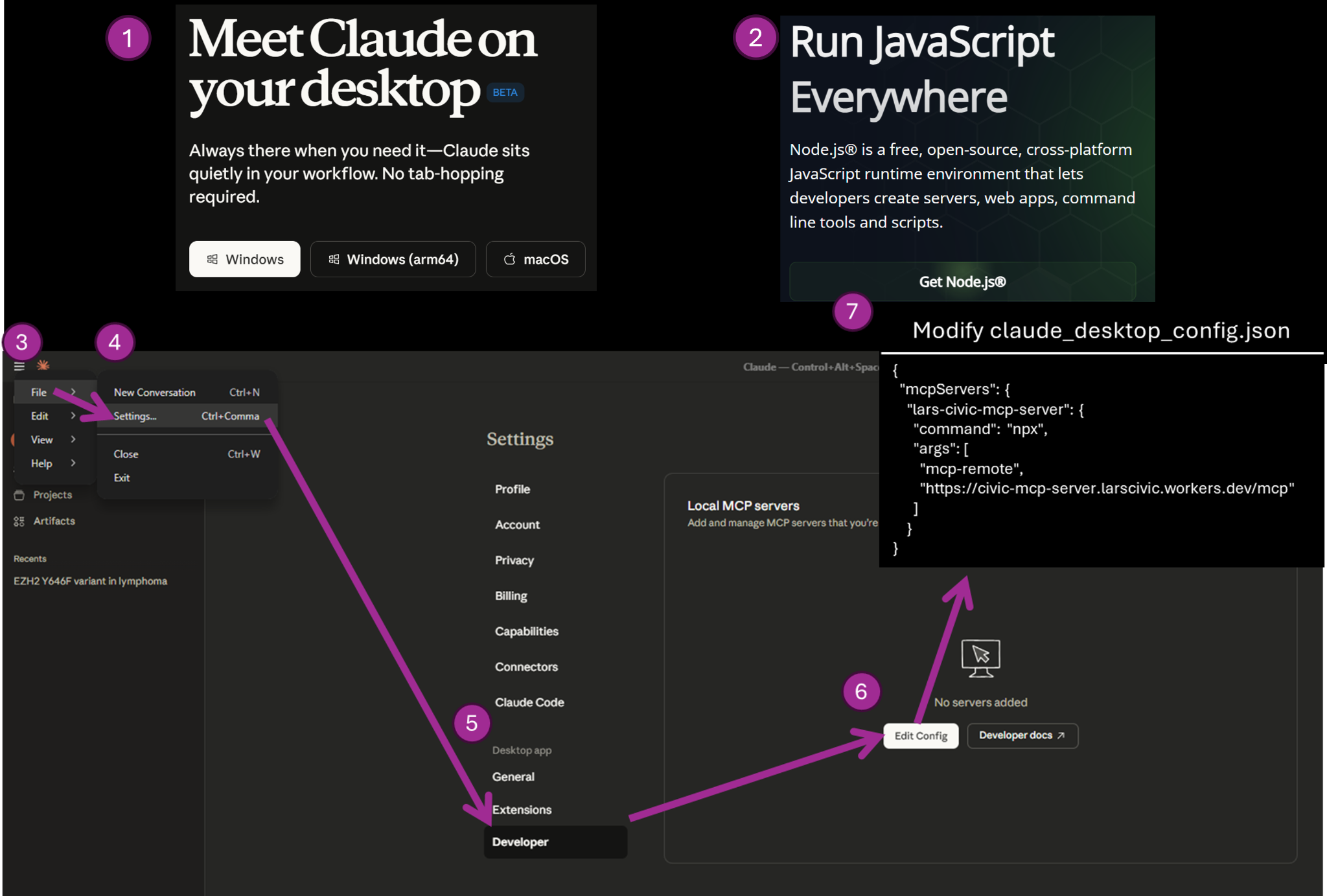

**Figure 1.** To interface with the CIViC MCP server through Claude Desktop, first install the application at <https://claude.ai/download>. Then install Node.js (LTS) from [https://nodejs.org](https://nodejs.org/en). After modifying claude_desktop_config.json, restart Claude Desktop. Claude will now have access to the CIViC MCP server during chats.

**Prompt 1.**

**System Prompt GPT.**

You are an expert biomedical annotator trained to assess variant evidence from the Clinical Interpretations of Variants in Cancer (CIViC) knowledgebase given a specific gene variant + cancer + therapy.

**System Prompt GPT + MCP.**

You are an expert biomedical annotator trained to assess variant evidence from the Clinical Interpretations of Variants in Cancer (CIViC) knowledgebase given a specific gene variant + cancer + therapy. Use the tools to answer oncology questions for the CIViC knowledgebase.

**User Prompt**

User: For this exact combination of gene variant: {variant}, cancer type: {cancer}, therapy: {therapy}, determine whether CIViC contains matching records for each evidence type + significance below, and whether those records support that significance.

Answer choices:

A. Supports

B. Does Not Support

C. No Evidence

CONTEXT-SENSITIVE EVIDENCE RULES (important):

- Treat the user's inputs (molecular profile, disease, therapy) as REQUIRED filters.

- Only count CIViC records that explicitly match ALL user-specified fields:

- Disease must match exactly.

- Therapy must match exactly as a set (order does not matter).

- Exception: if Therapy Interaction Type is SUBSTITUTES, overlap is sufficient only for single-drug queries; for multi-drug queries the full combination must match.

- Molecular Profile matching:

- If the QUERY MP contains "AND": only include evidence MPs that contain ALL components (no partial matches).

- If the QUERY MP contains "OR": include evidence MPs if ANY component matches.

- Otherwise (single MP): include evidence MPs that match that term; do NOT include evidence MPs with "AND".

TASK:

- For each item (<evidence type> — <significance>):

- Return A if there exists ≥1 matching record with evidenceDirection = SUPPORTS.

- Return B if there exists ≥1 matching record with evidenceDirection = DOES_NOT_SUPPORT.

- Return A,B if both SUPPORTS and DOES_NOT_SUPPORT exist for that item.

- If any matching record has evidenceDirection = NA / N/A / missing, treat direction as unknown and return A,B for that item.

- Return C only if there are zero matching records for that evidence type + significance.

- The significance label "N/A" refers ONLY to CIViC records whose significance field is NA or N/A (not UNCERTAIN_SIGNIFICANCE).

Output exactly one line per item below (23 lines), in this exact format:

<evidence type> — <significance>: <A|B|A,B|C>

Items (use exactly this order, and exactly these labels):

predictive — sensitivity_response: <A|B|A,B|C>

predictive — resistance: <A|B|A,B|C>

predictive — adverse_response: <A|B|A,B|C>

predictive — reduced_sensitivity: <A|B|A,B|C>

predictive — N/A: <A|B|A,B|C>

prognostic — better_outcome: <A|B|A,B|C>

prognostic — poor_outcome: <A|B|A,B|C>

prognostic — N/A: <A|B|A,B|C>

diagnostic — positive: <A|B|A,B|C>

diagnostic — negative: <A|B|A,B|C>

predisposing — predisposition: <A|B|A,B|C>

predisposing — protectiveness: <A|B|A,B|C>

predisposing — uncertain_significance: <A|B|A,B|C>

predisposing — N/A: <A|B|A,B|C>

oncogenic — oncogenicity: <A|B|A,B|C>

oncogenic — protectiveness: <A|B|A,B|C>

oncogenic — N/A: <A|B|A,B|C>

functional — gain_of_function: <A|B|A,B|C>

functional — loss_of_function: <A|B|A,B|C>

functional — unaltered_function: <A|B|A,B|C>

functional — neomorphic: <A|B|A,B|C>

functional — dominant_negative: <A|B|A,B|C>

functional — unknown: <A|B|A,B|C>

FINAL OUTPUT RULES:

- Output ONLY the 23 lines.

- No prose, no markdown, no explanations.

**Methods: Agent Mode Evaluation**

Because Agent Mode responses were substantially slower and subject to monthly usage limits, the benchmark was conducted across four paid ChatGPT accounts using a standardized protocol. To reduce user-specific bias, personalization features, including memory, were disabled for all accounts before testing. All evaluation prompts were conducted within a single project, with each prompt submitted in a fresh conversation window to minimize carryover from prior chat context. Response time was measured using the ChatGPT Timestamp Chrome extension, because the ChatGPT interface does not natively display message timestamps. Timing was defined as the interval from prompt submission to completion of the final Agent Mode response. To increase throughput, some prompts were run concurrently in separate tabs on the same account. This may have slightly inflated measured response times relative to strictly sequential single-session use. This consideration affects the timing comparison but not the classification outputs themselves.

**Table 1. GPT-5 + MCP Significance Results (Excluding “No Evidence”)**

| **Evidence Type** | **Significance** | **N** | **Precision** | **Recall** | **F1** |
| --- | --- | --- | --- | --- | --- |
| Predictive | Sensitivity | 28 | 1.00 | 0.96 | 0.98 |
| Predictive | Resistance | 17 | 1.00 | 0.94 | 0.97 |
| Predisposing | Predisposition | 13 | 1.00 | 1.00 | 1.00 |
| Oncogenic | Oncogenicity | 12 | 1.00 | 0.92 | 0.96 |
| Predisposing | N/A | 9 | 1.00 | 0.89 | 0.94 |
| Diagnostic | Positive | 8 | 1.00 | 1.00 | 1.00 |
| Predisposing | Uncertain Significance | 7 | 1.00 | 0.86 | 0.92 |
| Functional | Loss of Function | 6 | 1.00 | 1.00 | 1.00 |
| Functional | Gain of Function | 5 | 1.00 | 1.00 | 1.00 |
| Prognostic | Better Outcome | 5 | 1.00 | 1.00 | 1.00 |
| Prognostic | Poor Outcome | 5 | 1.00 | 1.00 | 1.00 |
| Functional | Dominant Negative | 4 | 1.00 | 1.00 | 1.00 |
| Predictive | Reduced Sensitivity | 3 | 1.00 | 1.00 | 1.00 |
| Diagnostic | Negative | 2 | 1.00 | 0.50 | 0.66 |
| Functional | Unaltered Function | 1 | 1.00 | 1.00 | 1.00 |

**Table 2. GPT-5 Significance Results (Excluding “No Evidence”)**

| **Evidence Type** | **Significance** | **N** | **Precision** | **Recall** | **F1** |
| --- | --- | --- | --- | --- | --- |
| Predictive | Sensitivity | 28 | 0.86 | 0.21 | 0.34 |
| Predictive | Resistance | 17 | 0.71 | 0.29 | 0.42 |
| Predisposing | Predisposition | 13 | 0.90 | 0.69 | 0.78 |
| Oncogenic | Oncogenicity | 12 | 0.25 | 0.17 | 0.2 |
| Predisposing | N/A | 9 | 0.00 | 0.00 | 0.00 |
| Diagnostic | Positive | 8 | 1.00 | 0.75 | 0.86 |
| Predisposing | Uncertain Significance | 7 | 0.00 | 0.00 | 0.00 |
| Functional | Loss of Function | 6 | 0.60 | 0.50 | 0.55 |
| Functional | Gain of Function | 5 | 0.75 | 0.60 | 0.67 |
| Prognostic | Better Outcome | 5 | 1.00 | 0.60 | 0.75 |
| Prognostic | Poor Outcome | 5 | 0.60 | 0.60 | 0.60 |
| Functional | Dominant Negative | 4 | 1.00 | 0.50 | 0.67 |
| Predictive | Reduced Sensitivity | 3 | 0.33 | 0.33 | 0.33 |
| Diagnostic | Negative | 2 | 0.00 | 0.00 | 0.00 |
| Functional | Unaltered Function | 1 | 0.00 | 0.00 | 0.00 |

**Table 3. GPT-5 Agent Mode Significance Results (Excluding “No Evidence”)**

| **Evidence Type** | **Significance** | **N** | **Precision** | **Recall** | **F1** |
| --- | --- | --- | --- | --- | --- |
| Predictive | Sensitivity | 28 | 1.00 | 0.79 | 0.88 |
| Predictive | Resistance | 17 | 1.00 | 0.88 | 0.94 |
| Predisposing | Predisposition | 13 | 0.93 | 1.00 | 0.96 |
| Oncogenic | Oncogenicity | 12 | 0.92 | 1.00 | 0.96 |
| Predisposing | N/A | 9 | 1.00 | 0.78 | 0.88 |
| Diagnostic | Positive | 8 | 0.89 | 1.00 | 0.94 |
| Predisposing | Uncertain Significance | 7 | 1.00 | 0.86 | 0.92 |
| Functional | Loss of Function | 6 | 0.86 | 1.00 | 0.92 |
| Functional | Gain of Function | 5 | 0.75 | 0.60 | 0.67 |
| Prognostic | Better Outcome | 5 | 1.00 | 1.00 | 1.00 |
| Prognostic | Poor Outcome | 5 | 1.00 | 0.80 | 0.89 |
| Functional | Dominant Negative | 4 | 1.00 | 1.00 | 1.00 |
| Predictive | Reduced Sensitivity | 3 | 1.00 | 1.00 | 1.00 |
| Diagnostic | Negative | 2 | 0.00 | 0.00 | 0.00 |
| Functional | Unaltered Function | 1 | 1.00 | 1.00 | 1.00 |
